# Planktonic diatom assemblage seasonal diversity used to assess environmental health in a coastal wetland of southern Chile

**DOI:** 10.1101/2020.08.19.257626

**Authors:** Catalina Ríos-Henríquez, Norka Fuentes, Claudio N. Tobar, Jaime R. Rau, Fabiola Cruces

**Affiliations:** Laboratorio de Limnología, Departamento de Acuicultura y Recursos Agroalimentarios, Universidad de Los Lagos, Av. Fuschlocher 1305, Osorno, Chile; Departamento de Ciencias Básicas, Universidad Santo Tomás, Los Carrera 753, Osorno, Chile; Laboratorio de Ecología, Departamento de Ciencias Biológicas & Biodiversidad, Universidad de Los Lagos, Campus Osorno, Casilla 933, Osorno, Chile; Departamento de Botánica, Facultad de Ciencias Naturales y Oceanográficas, Universidad de Concepción, Concepción, Chile

**Keywords:** diatoms, ecological diversities, plankton, coastal wetlands, river mouthd

## Abstract

Although planktonic diatoms are one of the most abundant taxonomic groups in coastal wetlands, their assemblages have not been used to determine the environmental health of these ecosystems. Studies of ecosystem environmental health have been based on other taxonomic groups; we propose that diatom genera diversity represents a viable alternative for this purpose. Thus, our aim was to determine the alpha and beta diversities of the planktonic diatom assemblage present in Caulín Bay, Chiloé Island (41° 49’S; 73° 38’O), southern Chile, during the austral winter and spring of the years 2012 and 2014. Inasmuch Caulín Bay is an important site for aquatic bird observation and conservation, hunting is prohibited on a national scale and, internationally, the site has been declared an Important Bird and Biodiversity Area (IBA). Our results indicate different diversities between sampling stations, but not between the years studied. In total, we recorded 53 diatom genera, of which the most abundant were *Coscinodiscus* (21.99%) and *Cocconeis* (16.23%). The study area presented high genera diversity (i.e., H’_(log2)_ >3.74) and beta diversity indicated that Caulín presents a low level of heterogeneity and is a low genera replacement environment. Consequently, we infer that Caulín Bay is productive and environmentally stable ecosystem. This leads us to conclude that diatom diversity determination is a viable alternative to establish aquatic ecosystem environmental health and we recommend that future conservation strategies be established for Caulín Bay.

**RESUMEN:** Si bien las diatomeas planctónicas son uno de los grupos taxonómicos más abundantes de los humedales costeros, sus ensambles no se han utilizado para determinar el estado ambiental de estos ecosistemas. Aunque se han realizado estudios de la salud ambiental de un ecosistema utilizando otros grupos taxonómicos, nosotros proponemos que la diversidad de géneros de diatomeas representa una alternativa viable. Por lo tanto, el objetivo de este estudio fue determinar las diversidades alfa y beta del ensamble de diatomeas planctónicas presentes en la Bahía de Caulín, Isla de Chiloé (41 ° 49’S; 73 ° 38’O), sur de Chile, durante las temporadas de invierno y primavera austral de los años 2012 y 2014. Bahía Caulín es un sitio importante para la observación y conservación de aves acuáticas por lo que a nivel nacional se ha prohibido la caza y a nivel internacional fue decretada un Área Importante para la Conservación de Aves, AICA. Los resultados de este estudio indicaron diferencias en las diversidades entre las estaciones de muestreo, pero no entre los años estudiados. En total, se identificaron 53 géneros de diatomeas; los más abundantes fueron *Coscinodiscus* (21,99%) y *Cocconeis* (16,23%). El área de estudio presentó una alta diversidad de géneros (i.e., H’_(log2)_ >3.74) y la diversidad beta indicó que Caulín presentó bajo nivel de heterogeneidad y es un entorno con bajo reemplazo de géneros. Así, inferimos que Bahía Caulín es un ecosistema productivo y ambientalmente estable, por lo cual concluimos que la determinación de las diversidades de diatomeas es una alternativa para establecer la salud ambiental de los ecosistemas acuáticos y recomendamos establecer futuras estrategias de conservación para Bahía Caulín.

PALABRAS CLAVE: diatomeas, diversidades ecológicas, plancton, humedales costeros, desembocadura

## INTRODUCTION

The environmental health of an ecosystem is usually assessed and characterized through species richness estimation, which provide information on the main biological diversity components (Nabout *et al*. 2007, Nogueira *et al*. 2008, Schuster *et al*. 2015). Spatial-temporal variability in communities is related to high species richness and, consequently, high productivity in the location (De Raedt *et al*. 2017). A current approach for determining the environmental health of a particular ecosystem is to establish the alpha and beta diversities of the primary producers’ taxa. In particular, alpha diversity is used to determine an ecosystem’s environmental stability, i.e., greater diversity implies more environmental stability, while beta diversity is used to determine ecosystem productivity, since high beta diversity values may indicate a productive environment (Siqueiros-Beltrones 2005; Zorzal-Almeida *et al*. 2017).

In aquatic ecosystems, diatoms play an important role within the trophic web (Agirbas *et al*. 2017), presenting high species diversity (Hutchinson 1967) especially in coastal zones (Iriarte *et al*. 2012). Consequently, they are considered to be indicators of environmental quality, mainly because they are present throughout the entire year, have short life cycles and are ecologically more easily recognized (Round *et al*. 1990).

Diatoms dominate in upwelling events during spring-summer when increased wind speeds transport nutrients from the bottom. They also dominate in winter, due to the contribution of continental subsidies rich in silica from rivers (González *et al*. 2007, Iriarte *et al*. 2010, Sánchez *et al*. 2012; Anabalón *et al*. 2016), which produce spatial-temporal variability and, as a result, changes in the environmental stability of ecosystems (Doherty *et al*. 2017). A study carried out in the Netherlands suggests there is a strong relationship between ecosystem environmental stability and diatom community beta diversity; that is, if the environment is eutrophic, the beta index decreases, so the turnover rate is negatively affected and environmental productivity decreases (Goldenberg *et al*. 2014).

In Chile, specifically the southern-austral coastal zone, diatoms contribute between 25 to 75% of primary productivity (Ochoa *et al*. 2010). Marine and freshwater diatoms are mixed (Avaria 2006), with intra-annual fluctuations in diversity due to seasonal variations in temperature, precipitation, water flow and upwelling (González *et al*. 2010). As a result, these fjord and channel ecosystems are of considerable ecological interest (Avaria *et al*. 1997). In particular, the high productivity in the inner sea of Chiloé Island is principally due to the contribution of continental subsidies and, to a lesser degree, upwellings (Iriarte *et al*. 2010).

Using diatom assemblages as predictors of ecosystem environmental health could be a viable method, due to their limited dispersal potential, short generation period, and because they are sensitive to changes in nutrient concentrations of anthropic origin. The latter would be reflected in variations in their composition and abundance on a temporal and spatial scale (Lai *et al*. 2018; Kaufouris *et al*. 2019) which, in turn cause differential changes in diversity, depending on the type and level of contamination (Stevenson *et al*. 2010; Pandey *et al*. 2018). In addition, taxonomic identification to genus level is sufficient for use as eutrophication indicators, or to establish assemblage patterns for studies related to environmental conservation (Heino & Soininen 2007; Kaufouris *et al*. 2019). However, knowledge about the environmental health of aquatic ecosystems based on diatom community studies is scarce (Goldenberg *et al*. 2014, Zorzal-Almeida *et al*. 2017) and in Chile none reported so far. Therefore, the goal of this study was to characterize the environmental health of Caulín Bay coastal wetland by estimating diatom genera richness. To this end we determined the alpha and beta ecological diversities of the diatom assemblages during the years 2012 and 2014. The information provides on the principal ecological diversity components in relation to environmental variables, both seasonally and annually, necessary to establish effective conservation measures in the area.

## MATERIALS AND METHODS

### STUDY AREA

Caulín Bay (41°49’S; 73°38’W) is located in the northern zone of Chiloé Island, southern Chile (Figs. 1a, 1b) and has a maximum depth of 5 m. The Huenque river empties into Caulín Bay at a flow rate that fluctuates from 8.92 m^3^/s in the austral winter to 1.60 m^3^/s in the austral spring (Ríos 2015). The climate is temperate maritime, with high rainfall and Mediterranean influence. Precipitation ranges between 1.000 and 3.000 mm, with highest rainfall in the months of May and August (>50% of pp) (Delgado *et al*. 2010). Average temperature during the austral winter and spring is 5.7°C and 11.3°C respectively (METEOCHILE 2018). Caulín Bay is considered an important site for aquatic bird observation and conservation both nationally (i.e., hunting is prohibited) and internationally (i.e., it is an IBA site (Important Bird and Biodiversity Area) (Cursach *et al*. 2015) and has recently been declared “Espacio Marítimo Costero de Pueblos Originarios” (EMCPO) (Marine and Coastal Area for Indigenous Peoples (SUBPESCA 2018).

**Figure 1:**
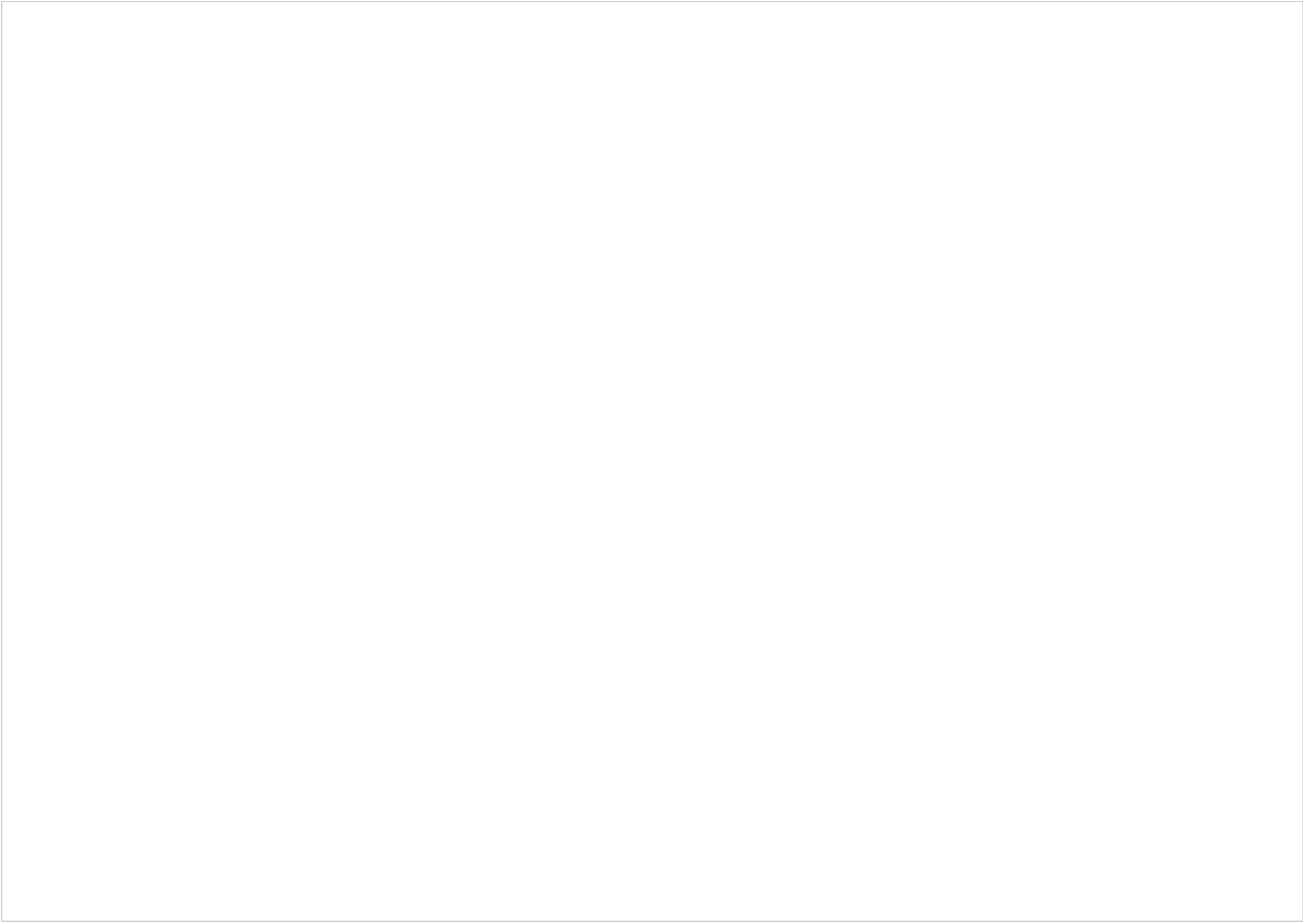
Geographic location of the study area, Caulín Bay (41°49’S; 73°38’W). A: a map of Chile, the black square indicates Chiloé Island; B: a map of Chiloé Island, highlighting the location of Caulín Bay; C: the location of the sampling points (the grey circles were only sampled in 2014). BC: Caulín Bay; RH: Huenque River. / Localización geográfica del área de estudio, Bahía de Caulín (41°49‥S; 73°38′O). A: muestra un mapa de Chile, el cuadrado negro indica a la isla de Chiloé; B: muestra un mapa de la isla de Chiloé, señalando la localización de la Bahía de Caulín; C: muestra la localización de los puntos de muestreo (los círculos grises sólo fueron muestreados en 2014). BC: Bahía de Caulín: RH: Río Huenque.

### SAMPLING

We collected planktonic diatom samples during the austral winter and spring of 2012 and 2014, both years with a neutral ENSO (El Niño-Southern Oscillation) (METEOCHILE 2018). We established nine sampling points along the length and width of Caulín Bay and two at the mouth of the Huenque river. In 2014 we reduced the sampling points to five (3 in the bay and 2 at the river mouth) (Fig. 1c), given that no significant taxa spatial variation was recorded in the 2012 samples (ANOSIM; p = 0.1). We collected samples by horizontal hauling over 10 m stretches using a plankton net with 30 µm mesh size filtering 0.71 m^3^ of water samples were placed in dark 500 ml containers and fixed with Lugol (Hötzel & Croome 1999). We established taxonomic determination to genus level and abundance (cells ml^-1^) with a LEITZ DIAVERT inverted microscope. We adopted the Utermöhl (1958) methodology, using a 25 ml sedimentation chamber where we established 2 transects (transversal and longitudinal), with the aid of specialized literature to identify diatoms at the genera level of taxonomic resolution (Rivera & Valdebenito 1979, Hasle & Syvertsen 1997, Inostroza *et al*. 2010).

We refer to Caulín Bay as BC; Huenque river as RH; austral winter (W) or austral spring (S), in relation to the season of the sample, and finalize with the corresponding year (12 or 14) (e.g., BCW12).

### STATISTICAL ANALYSIS

We determined alpha diversity through genera richness, after determining sampling effort with Rarefaction estimators, expected genera richness with non-parametric Chao 1 and Jackknife estimators which were obtained using the PAST statistical programs and Krebs (1999) ad hoc computation program (EcoMeth software). We also calculated the Shannon index of diversity (H’_(log2)_), with its respective variance, using the DIVERS program in the FRANJA computational package (Franja 1993). We then compared these results with the Hutcheson (1970) statistical test for p=0.05. Given that, in theory, H’ can vary from zero to very high values (cf. Krebs 1979) but, in practice, it becomes asymptotic at values of five bits/taxon, we considered high alpha diversity as H’>5 bits/taxon (see Rau *et al*. 1998). We regarded normal or moderately high diversity as ranging between H′=2.6 - 3.8 values (see Siqueiros-Beltrones *et al*. 2017).

Whittaker (1972) standardized beta diversity (Dβ) was obtained from Rau *et al*. (1998). Values vary between zero and one, where zero indicates an identical assemblage and one, a different assemblage.

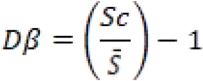

Where:

S_c_= genera richness in a composite sample (i.e., which combines a given number of paired α samples).

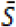= average (arithmetic mean) genera richness in the paired α samples compared

We also used another beta diversity (β), proposed by Schluter & Ricklefs (1993) based on the formula obtained from Moreno (2001):

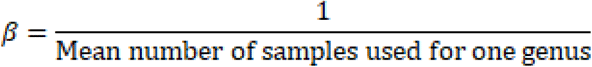

Subsequently, we constructed an ecological matrix of genera abundance per site by carrying out square root transformations, and calculated the Bray-Curtis dissimilitude matrix. We carried out similarity analysis (one-way ANOSIM) to analyze differences between years and used the Mann-Whitney U (α=0.05) test to identify spatial differences between the river and the bay during each season. We employed a Similarity Percentage Analysis (SIMPER) to identify which taxa contributed to the significant differences between assemblages. We used non-parametric statistics when data did not meet the assumptions of normality and homoscedasticity required in the current parametric statistics. We conducted the aforementioned analyses with the PRIMER v6 statistical program (Clarke & Gorley 2006) and VassarStats: Website for Statistical Computation (<http://www.vassarstats.net>), the latter was used for Mann-Whitney U test. Finally, using the same software, we applied a Spearman’s non-parametric correlation test to measure similitude between Whittaker (1972) and Schluter & Ricklefs (1993) beta diversity indexes.

## RESULTS

Based on genera richness we identified a total of 53 diatom genera: 33 and 32 recorded during the austral winter and spring of 2012; and 30 and 40 during the austral winter and spring of 2014, where 60.71% corresponded to the Class Bacillariophyceae; 21.43% to the Class Mediophyceae and 17.86 % to the Class Coscinodiscophyceae. In 2012, the most abundant genera in the austral winter were *Odontella* (12.18 %) and *Thalassiosira* (7.22 %) and in the austral spring, *Navicula* (21.33 %) and *Campylosira* (12.59 %). In 2014 *Coscinodiscus* (59.94 %) and *Cocconeis* (12.41 %) were the most abundant genera in the austral winter, and *Cocconeis* (36.10 %) and *Navicula* (18.20 %) in the austral spring (Table 1).

**Table 1:**
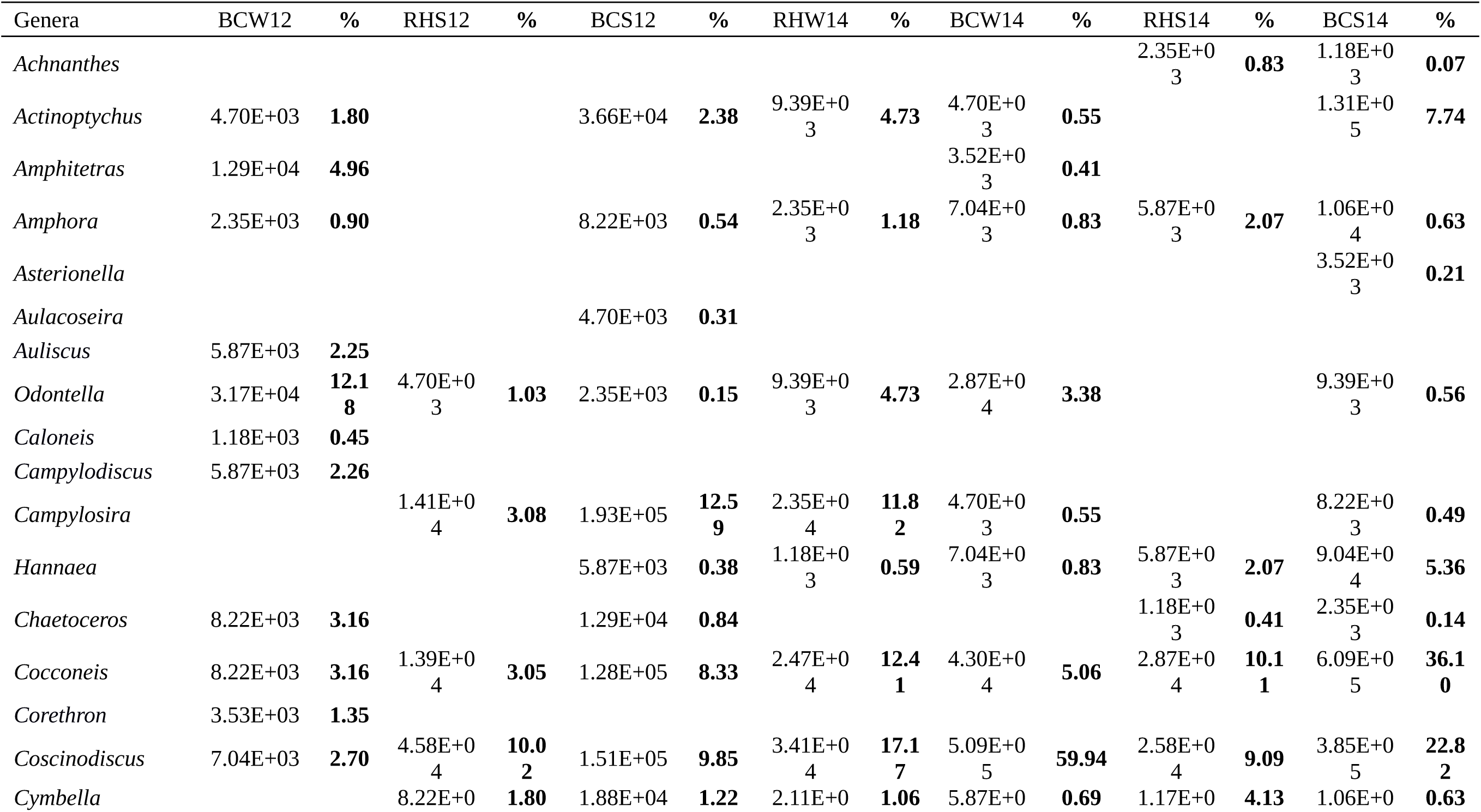
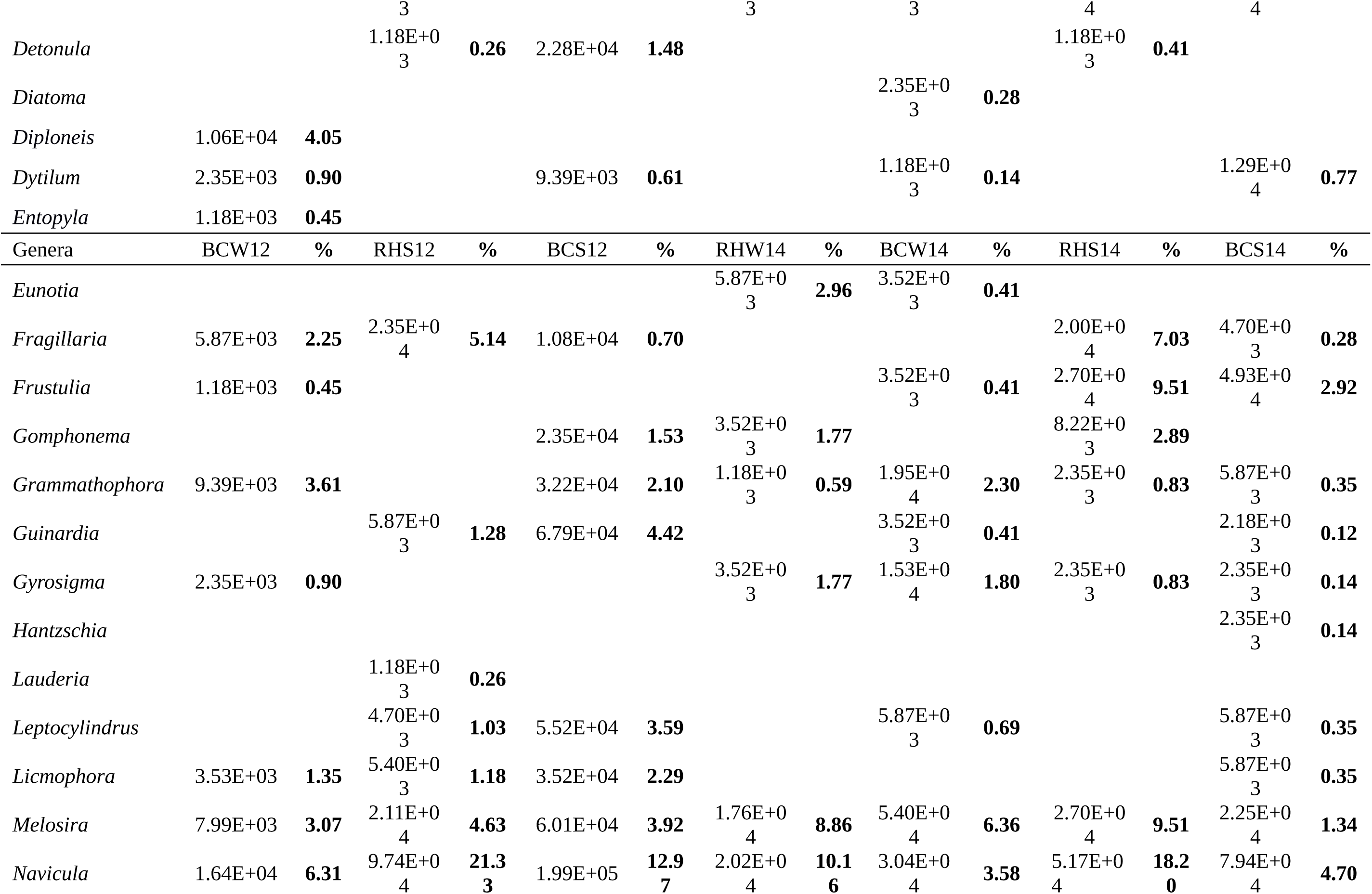
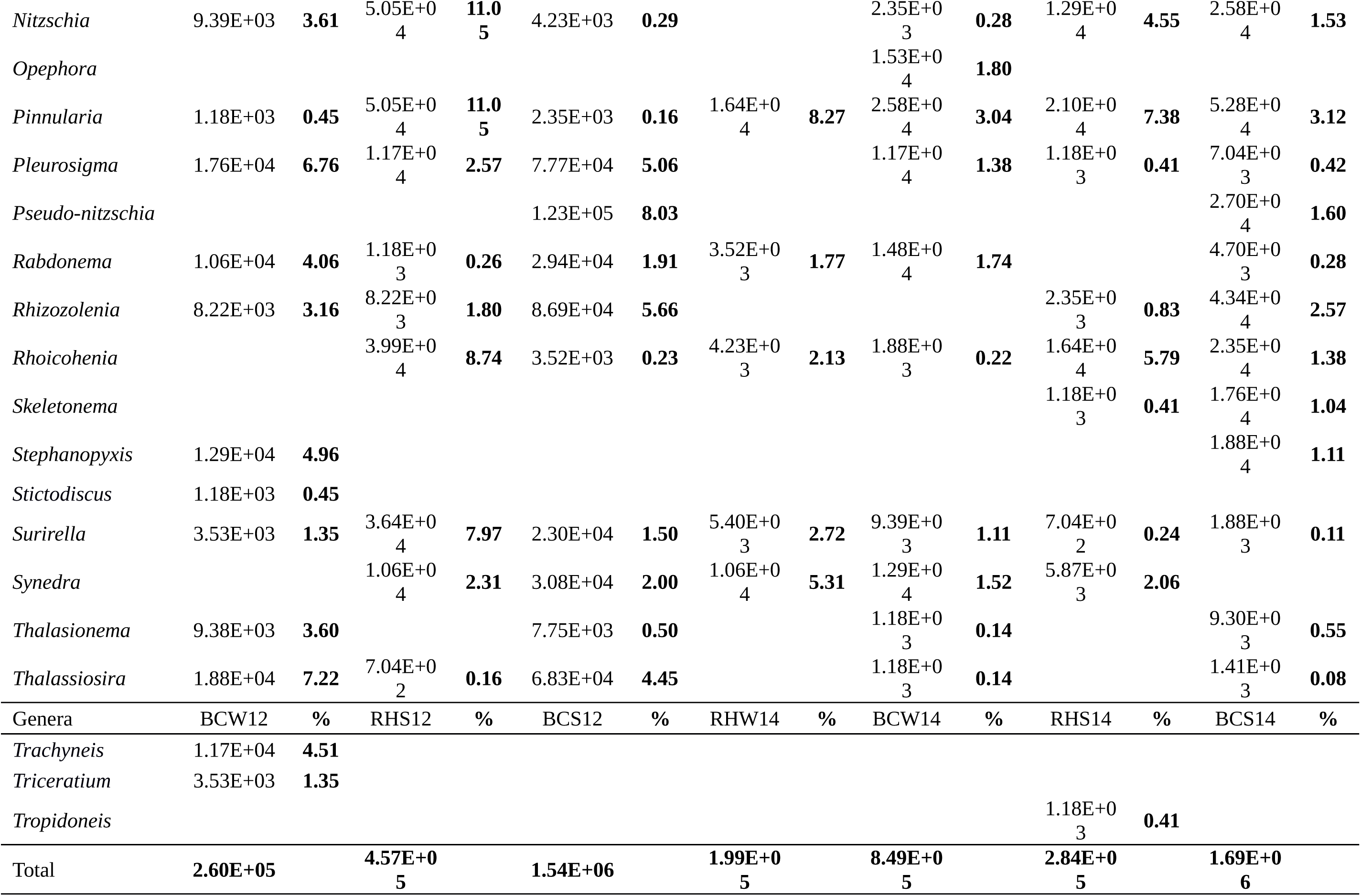
Abundance (cell m^-3^) of diatom genera identified during the study periods. We refer to Caulín Bay as BC; Huenque river as RH; austral winter (W) or austral spring (S), in relation to the season of the sample, and finalize with the corresponding year (12 or 14) (e.g., BCW12 or RHW14). / Abundancia (cell m^-3^) de géneros de diatomeas encontrados durante los períodos de estudio. Nos referimos a la Bahía de Caulín como BC; Río Huenque como RH; invierno austral (W) o primavera austral (S), en relación con la temporada de la muestra, y finalizar con el año correspondiente (12 o 14) (por ejemplo, BCW12 o RHW14).

Alpha diversity was generally high (i.e., H’(log_2_) >3.74), we recorded the highest value in the bay, specifically in the BCW12 station (H’(log_2_) =4.58). The Hutchenson analysis showed that the winter of 2012 presented significant statistical differences with the majority of the sampling stations (p<0.05) (Table 2). Furthermore, although we recorded differences between the austral winter and spring (p=8.71×10-21), no differences were found between the river and bay.

**Table 2:** Hutcheson test results for comparing diversities, those with significant (p<0.05) differences in grey. We refer to Caulín Bay as BC; Huenque river as RH; austral winter (W) or austral spring (S), in relation to the season of the sample, and finalize with the corresponding year (12 or 14) (e.g., BCW12 or RHW14). / Resultados de la prueba de Hutcheson para comparar diversidades, aquellos con diferencias significativas (p<0,05) se señalan en gris. Nos referimos a la Bahía de Caulín como BC; Río Huenque como RH; invierno austral (W) o primavera austral (S), en relación con la temporada de la muestra, y finalizar con el año correspondiente (12 o 14) (por ejemplo, BCW12 o RHW14).

| Samples | BCS12 | RHW1<br>4 | BCW14 | RHS14 | BCS14 |
| --- | --- | --- | --- | --- | --- |
| BCW12 | 9.20x10<sup>-7</sup> |  | 2.59 x10<sup>-9</sup> |  |  |
| RHS12 |  | 0.92 |  | 0.66 |  |
| BCS12 |  |  | 0.005 |  | 3.38 x10<sup>-6</sup> |
| RHW14 |  |  |  | 0.62 |  |
| BCW14 |  |  |  |  | 0.41 |
| RHS14 |  |  |  |  |  |
| BCS14 |  |  |  |  |  |

According to the rarefaction curve and the Chao 1 non-parametric estimator, sampling effort and expected genera richness were sufficient to reveal a diatom genera richness representative of the sites, with the exception of site BCW12. In the latter case, the curve did not reach the asymptote (200 individuals, 20 genera), due to differences in the count in river a minimum of 250 individuals was required and in bay >500 individuals (Fig. 2). Furthermore, richness observed was similar to the richness obtained through the Chao 1 non-parametric estimator (Table 3). We recorded 20 unique genera in site RHS14 using a Jackknife procedure (Krebs 1989), while only two unique genera were identified in site BCS12. Similarly, Jackknife analysis indicated highest estimated richness (35 genera) in site BCW12, and lowest estimated richness (25 genera) in site RHW14; in the BCS12 site observed richness coincided with estimated richness (31 genera).

**Table 3:** Alpha diversity and diatom genera richness in the study area. We refer to Caulín Bay as BC; Huenque river as RH; austral winter (W) or austral spring (S), in relation to the season of the sample, and finalize with the corresponding year (12 or 14) (e.g., BCW12 or RHW14). / Diversidad alfa y riqueza de géneros de diatomeas presentes en el área de estudio. Nos referimos a la Bahía de Caulín como BC; Río Huenque como RH; invierno austral (W) o primavera austral (S), en relación con la temporada de la muestra, y finalizar con el año correspondiente (12 o 14) (por ejemplo, BCW12 o RHW14).

|  | H' (log 2) |  | Jackknife | Chao 1 |  |
| --- | --- | --- | --- | --- | --- |
| Samples | Index | Observed Richness | Estimated genus Richness | Number of unique genus | Index |
| BCW12 | 4.58±0.002 | 33 | 34.6±2.71 | 6 | 35.5 |
| RHS12 | 3.79±0.002 | 22 | 25.5±1.5 | 7 | 25 |
| BCS12 | 4.12±0.000<br>4 | 31 | 32.8±1.18 | 2 | 31 |
| RHW14 | 3.79±0.002 | 19 | 25±0.5 | 7 | 19.5 |
| BCW14 | 3.98±0.002 | 29 | 33±2.31 | 6 | 30 |
| RHS14 | 3.74±0.002 | 25 | 34.3±5.5 | 20 | 27 |
| BCS14 | 3.91±0.001 | 34 | 40±1.15 | 8 | 34 |

**Figure 2:**
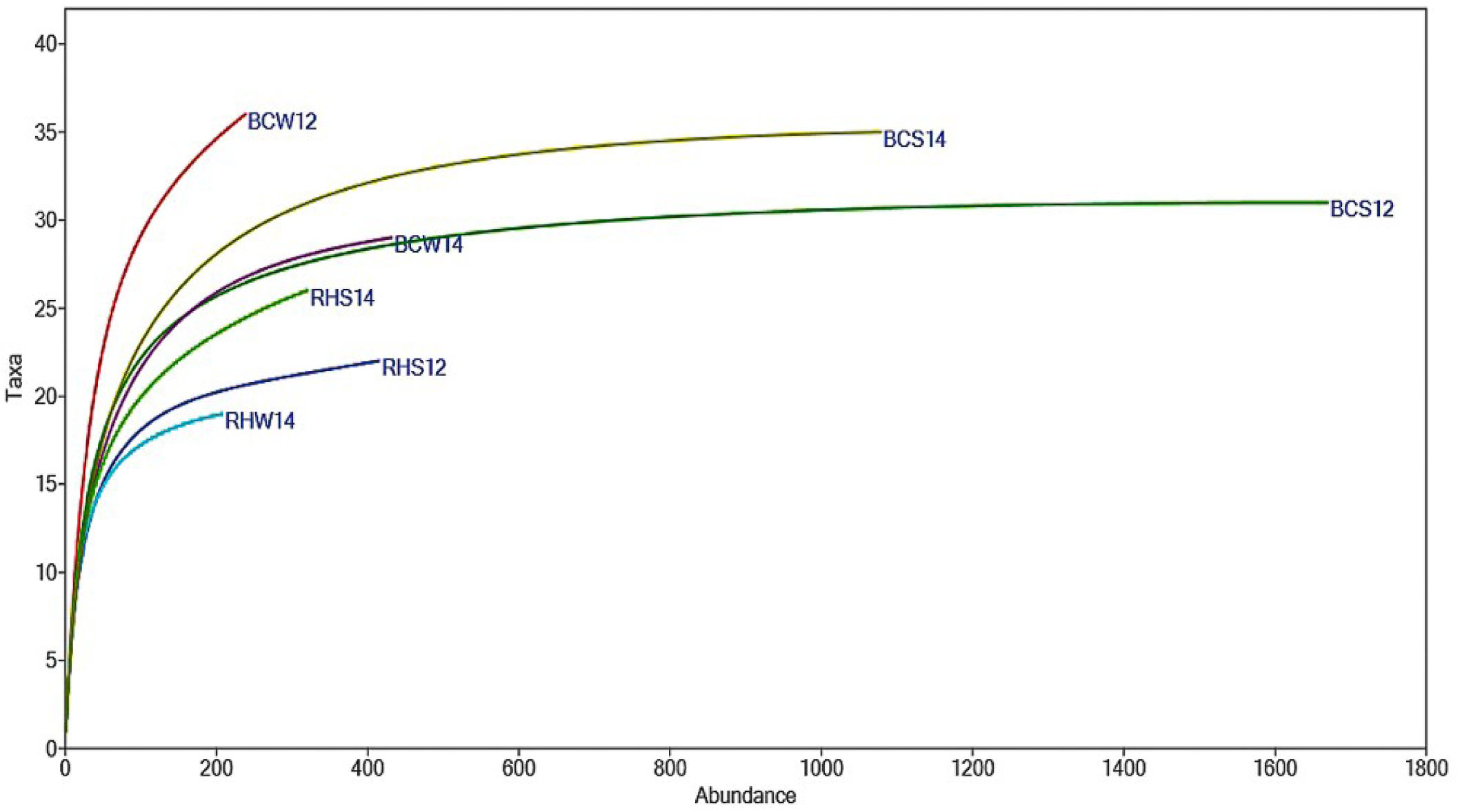
Rarefaction curve over the period we studied diatom genera. S: austral spring; W: austral winter. We refer to Caulín Bay as BC; Huenque river as RH; austral winter (W) or austral spring (S), in relation to the season of the sample, and finalize with the corresponding year (12 or 14) (e.g., BCW12 or RHW14). / Curva de rarefacción para el período de tiempo que los géneros de diatomeas fueron estudiados. S: primavera austral; W: invierno austral. Nos referimos a la Bahía de Caulín como BC; Río Huenque como RH; invierno austral (W) o primavera austral (S), en relación con la temporada de la muestra, y finalizar con el año correspondiente (12 o 14) (por ejemplo, BCW12 o RHW14).

Whittaker’s beta diversity (1972) rendered a value of Dβ=0.31, with the highest value in site RHS14 (Dβ=0.73) and the lowest in site RHS12 (Dβ=0.19). The beta diversity index proposed by Schluter & Ricklefs (1993) produced values of β≈0.6, where the highest value was in site RHS14 (β=0.87) and the lowest in the BCS12 (β=0.19) (Table 4). The correlation (Spearman’s non-parametric test) between the beta diversities of Whittaker (1972) and Schluter & Ricklefs (1993) was negative (rs = -0.314), but we were not able to obtain P value due the small sample size (n = 6).

**Table 4:** Beta diversity (Dβ) results from the study sites (CBW12 is excluded, no replicates in the river). We refer to Caulín Bay as BC; Huenque river as RH; austral winter (W) or austral spring (S), in relation to the season of the sample, and finalize with the corresponding year (12 or 14) (e.g., BCW12 or RHW14). / Resultados de diversidad (Dβ) para los sitios de estudio (CBW12 se excluye; no hubo réplicas para el río). Nos referimos a la Bahía de Caulín como BC; Río Huenque como RH; invierno austral (W) o primavera austral (S), en relación con la temporada de la muestra, y finalizar con el año correspondiente (12 o 14) (por ejemplo, BCW12 o RHW14).

| Samples | Whittaker (1972) | Schluter & Ricklefs (1993) |
| --- | --- | --- |
| RHS12 | 0.19 | 0.59 |
| BCS12 | 0.71 | 0.19 |
| RHW14 | 0.23 | 0.61 |
| BCW14 | 0.32 | 0.44 |
| RHS14 | 0.70 | 0.87 |
| BCS14 | 0.46 | 0.49 |

Diatom temporal abundance did not differ significantly either between the austral winters of 2012 and 2014, or between the austral springs of 2012 and 2014. Similarly, we did not record any differences between the austral winter and austral spring seasons of the years 2012 and 2014 (ANOSIM; R (Global) =1; p=0.333). However, the Mann-Whitney U test showed statistically significant differences in spatial abundance reported in the austral springs of 2012 and 2014, between RH and BC ecosystems (p>0.05). Only the austral winter of 2014 did not present statistically significant differences (p<0.05) between the two locations.

According to the SIMPER analysis, the genera that influenced spatial variability in the bay were a) between the austral winter and spring of 2012: *Campylosira* and *Coscinodiscus* (7.52 and 7.04%, respectively), we only recorded *Campylosira* in the austral spring; b) between the austral spring of 2012 and the austral spring of 2014: *Campylosira* and *Pseudo-nitzschia* (7.87 and 5.07%, respectively), these genera were more abundant in the austral spring of 2012, decreasing in 2014; c) between the austral winter and spring of 2014: *Actinoptychus* and *Coscinodiscus* (7.77 y 7.56%, respectively), greatest abundances were in the austral spring; and finally d) between the austral winter of 2012 and winter of 2014: *Coscinodiscus* and *Cocconeis* (7.29 y 4.23 %, respectively), they were more abundant in 2014.

Taxa that presented dissimilarity in the river were *Frustulia* and *Campylosira*, between the austral spring of 2012 and the austral spring of 2014 (8.33 and 6.01%, respectively) and between the austral winter of 2014 and the austral spring of 2014 (9.04 and 8.43%, respectively). We also identified the genera that contributed to dissimilarity between seasons: we did not record *Campylosira* in the austral winter of 2012, but we did identify its presence in the austral winter of 2014; we recorded *Navicula* in the austral spring of 2014, but not in the austral spring of 2012. We also observed genera differences in 2012 recording *Guinardia* in the austral spring, but not in the austral winter. On the contrary, we recorded *Amphitetras* and *Stephanopyxis* in the austral winter of the same year, but not in the austral spring. In 2014, we only identified *Navicula* and *Fragilaria* in the austral spring, while *Eunotia* and *Opephora* were only present in the austral winter (Table 1). However, similarities did exist between RH and BC in the austral winter of 2014 (63.30%) and between sites BCS12 and BCS14 (59.26%). On the other hand, shows that site BCW12 was only similar in 39.35% to the other sample sites (Pi = 3.98; p = 0.001), the remainder being significantly different.

## DISCUSSION

Determination of alpha and beta diversity using diatom taxa was useful to establish ecosystem health in Caulin Bay and Huenque river. The richness and diversity of diatom genera recorded in both Caulín Bay and at the Huenque river mouth was high, mainly because the study area is a semi-enclosed bay less than 5 m depth. According to Prado (1996) and Goldenberg *et al*. (2014) habitat size influences positively in diversity and richness patterns, particularly in the case of shallow, semi-closed coastal areas, since they limit oceanic currents access, thus avoiding high diatom dispersion, i.e., the residence period is longer, they are able to reflect the negative or positive impacts on the environmental health of the ecosystem (Hillebrand and Kahlert 2001, Kafouris *et al*. 2019). The significant diatom diversity and richness in Caulin Bay is mainly due, among other factors (e.g., grazing, pH or salinity) to the contribution of continental silica from the Huenque river and other smaller tributaries (Ríos 2015). It is well known that rivers are silica contributors of in coastal areas (Trigueros & Orive 2001, Cloern *et al*. 2017).

The alpha diversity obtained in this study, according to the rarefaction curve, Chao 1 non-parametric estimator and Jackknife procedure, indicates that the sampling effort was sufficient to ensure that results obtained were representative of the site; that is, representative of the planktonic diatom genera richness present in the assemblage. In conservation studies it is important to determine the sampling effort, which represents species richness in the location the greater the sampling effort, the closer the richness value is the ecosystem reality. It also indicates that diversity patterns are consistent (Gaston 1996, Schuster *et al*. 2015). In our study, *Cocconeis* and *Coscinodiscus* were the dominant genera, contrary to more northern zones of Chile where the dominant genera were *Rhizosolenia, Chaetoceros* and *Nitzschia*. These genera are typically found in the Peru current (Prado 1996), associated with warmer temperatures (17° to 20° C), although their diversity is low, as is their species abundance (Torres-Zambrano 1998). In the extreme south of Chile, specifically in the Antarctic zone, high dominance of *Thalassiosira, Corethron* and *Skeletonema* has been found, these genera are considered resistant to thermal anomalies (Torres-Zambrano & Tapia 1998, Paredes *et al*. 2014).

The proximity between observed and estimated genera richness shows that the sampling effort was sufficient to reveal diatom assemblage diversity (Nogueira *et al*. 2010, Schuster *et al*. 2015). For example, in the BCS12 sample observed richness was 31, Jackknife estimator was 32.8 ± 1.18 and Chao 1 estimator was 31 (Table 3). Although the Jackknife procedure is not commonly employed in microalgae studies (Nabout *et al*. 2007, Nogueira *et al*. 2008), it made a significant contribution to our study because it shows the number of unique genera recorded in each sample location (Table 3). These values reveal environmental heterogeneity, as opposed to the prevalence of common species over time, which indicates environmental homogeneity (Olden & Rooney, 2006). Therefore, according to alpha diversity, Caulín Bay is a heterogeneous, healthy environment.

We used the Whittaker (1972) and Schluter & Ricklefs (1993) indexes to determine beta diversity in Caulín Bay and results indicated statistical differences in the values obtained between sites. Thus, a higher value of 0.71, close to one, in the BCS12 site (bay) is indicative of a more heterogeneous, and therefore more productive, environment as compared to the RHS12 site (river), where the value obtained (0.19) is closer to zero, suggesting that this sampling station is homogeneous and, consequently, less productive (Zorzal-Almeida *et al*. 2017) (Table 4). In general, our results indicate a moderate beta diversity, that is, a low level of heterogeneity and still productive. However, this result reveals that the environmental health of Caulín Bay is beginning to deteriorate (Schuster *et al*. 2015). Recent studies, conducted by Zorzal-Almeida *et al*. (2017) identified a relationship between productivity and beta diversity, because natural and gradual enrichment can increase the beta diversity. However, an accelerated increase in productivity due to anthropogenic enrichment can lead to unfavorable conditions and, consequently, to a decrease in beta diversity (Donohue *et al*. 2009). This phenomenon has not yet occurred in the coastal wetland of Caulín Bay, as indicated by alpha diversity. On the other hand, beta diversity indicates a slight deterioration in environmental health, possibly due to small-scale cultivation of “pelillo” (*Gracilaria chilensis* Bird, McLachlan & Oliveira, 1986) in the bay, which is why we believe protection measures must be established in Caulín Bay in order to avoid greater intervention in this wetland, particularly since, at present, conditions to sustain the diversity associated with this coastal wetland appear stable.

The use of alpha and beta diversities to determine productivity and ecosystem health has been useful and transversal, given that studies have been conducted both in terrestrial (Harrison *et al*. 2006) and aquatic (Langenheder *et al*. 2012, Bini *et al*. 2014) ecosystems, studying different biological groups. This include benthic macroinvertebrates (Heino *et al*. 2013), phytoplankton (Mousing *et al*. 2016), macrophytes (Thomaz *et al*. 2003), terrestrial insects (Kadowaki & Inouye 2015) and vertebrates (Martin & Ferrer 2015). Our study based on diatom assemblages was useful to infer the ecosystem’s environmental health.

Regarding spatial variation in the Caulín Bay, we observed homogeneity in the genera identified in the BC and RH sampling sites. However, on comparing both ecosystems (BC and RH) according to the U de Mann-Whitney test analyses, they were spatially different during the austral spring, where salinity played a significant role in the genera difference recorded between these locations (Campanelli *et al*. 2009, Bode *et al*. 2017). No differences were recorded in the austral winter between RH and BC, due to the greater mix during this period, as a result of wind action and increased water flow (Zheng & DiGiacomo 2018). Since Caulín is a semi-enclosed bay, it presents homogeneous environmental conditions (Ríos 2015) and, accordingly, diatom spatial distribution would not vary significantly, although seasonal changes would occur, as we found in this study.

The temporal variation observed in our study indicates predominance of Coscinodiscophyceae diatoms during the austral winter, when high concentrations of silica input from rivers occur (Iriarte *et al*. 2010). This coincides with the study conducted by Labbé-Ibáñez *et al*. (2015) in the Seno of Reloncaví, southern Chile. In contrast, the austral spring we identified Coscinodiscophyceae, Mediophyceae and Bacillariophyceae. According to González *et al*. (2011) this is due to the greater mix between sea-water and river discharge during the austral spring. The Bacillariophyceae, in general, predominate in shallow environments, with low silica concentrations, while the Coscinodiscophyceae and Mediophyceae predominate when silica concentration is higher (Reynolds 2006).

Finally, we conclude that determination of alpha and beta diversities in diatom assemblage, as a case study, proved a useful method to verify the stability and productivity of the Caulín Bay coastal wetland. In spring, beta diversity was moderate in the bay, but low in the river, corroborating spatial variation (between river and bay). Diatom genera abundance and richness recorded in Caulín Bay were high, indicating this is a highly productive site where primary productivity is sufficient to maintain stable condition in this coastal wetland. In view of this, we recommend that conservation and management policies be established for the Caulín Bay coastal wetland. These measures would avoid a decrease in primary productivity (diatoms) which would negatively affect the higher trophic levels (grazers), since birds and other species depend directly or indirectly on these microalgae.

## ACKNOWLEDGEMENTS

The authors acknowledge the financial support of the Dirección de Investigación of the Universidad de Los Lagos, through Research Nucleus BIODES and BIODES 2.0. Our thanks also to Dr. Boris López for reviewing the manuscript and to Susan Angus for translating the text.

